# State-behavior feedbacks suppress personality variation in boldness during foraging in sticklebacks

**DOI:** 10.1101/2020.07.23.217828

**Authors:** Hannah E. A. MacGregor, Aislinn Cottage, Christos C. Ioannou

**Affiliations:** School of Biological Sciences, University of Bristol, Bristol, BS8 ITQ, U.K

**Keywords:** Animal temperament, risk-taking, repeatability, refuge use, *Gasterosteus aculeatus*, consistency

## Abstract

Consistent inter-individual variation within a population, widely referred to as personality variation, can be affected by environmental context. Feedbacks between an individual’s personality and state can strengthen (positive feedback) or weaken (negative feedback) individual differences when experiences such as predator encounters or winning contests are dependent on personality type. We examined the influence of foraging on individual-level consistency in refuge use (a measure of risk-taking, i.e. boldness) in three-spined sticklebacks, *Gasterosteus aculeatus*, and particularly whether changes in refuge use depended on boldness measured under control conditions. In the control treatment trials with no food, individuals were repeatable in refuge use across repeated trials, and this behavioral consistency did not differ between the start and end of these trials. In contrast, when food was available, individuals showed a higher degree of consistency in refuge use at the start of the trials versus controls but this consistency significantly reduced by the end of the trials. The effect of the opportunity to forage was dependent on personality, with bolder fish varying more in their refuge use between the start and the end of the feeding trials than shyer fish. This suggests a state-behavior feedback, but there was no overall trend in how individuals changed their behavior, with some individuals spending more, and others less, time in the refuge area at the end than at the start of the trials. Our study shows that personality variation can be suppressed in foraging contexts and a potential but unpredictable role of feedbacks between state and behavior.

## INTRODUCTION

Individuals of the same species within a population often differ consistently in their behavior over time and contexts (Dall et al., 2004; Magurran et al., 1998; Sih et al., 2004). For instance, individuals may be consistent in their reaction to environmental stimuli such as food (MacGregor et al., 2020; Szopa-Comley et al., 2020) or predators (Boissy, 1995), or express behavioral correlations across contexts such as being more aggressive to conspecifics and also bolder in the presence of predators (Huntingford, 1976). It is increasingly evident that this personality variation can be heritable (Oers et al., 2004), has fitness consequences (Smith and Blumstein, 2008), and contributes to a diverse range of ecological and evolutionary processes (Carere and Gherardi, 2013; Dall et al., 2012; Dingemanse and Réale, 2005).

Recently, attention has turned to understanding the conditions that promote the expression of animal personality, and the interplay between personality and behavioral plasticity (Briffa et al., 2008; Dingemanse et al., 2010; Mathot et al., 2011). On the one hand, plasticity in behavioral traits can allow individuals to respond rapidly and adaptively to changing conditions, including to factors such as predation risk, resources, and the social environment (Snell-Rood, 2013; Via et al., 1995). On the other hand, personality variation can arise under fluctuating selection where the optimal behavioral phenotype varies over space or time (Boon et al., 2007; Mangel, 1991). Conventionally, personality and plasticity in behavior have been studied independently, however it is increasingly apparent that they may co-vary, with individuals differing in their responsiveness to changes in environmental and social conditions (Bevan et al., 2018; Biro et al., 2010; Dingemanse et al., 2012; Laskowski and Bell, 2013; Stamps, 2016; Westneat et al., 2011).

The effects of environmental and social factors on the expression of behavioral variation may be linked to the state-dependence of behavior, where state variables refer to intrinsic factors (e.g. morphology, physiology, information, fecundity) that influence the balance between the costs and benefits of an animal’s behavioral decisions (Dingemanse and Wolf, 2010; Houston et al., 1999; Sih et al., 2015). It is widely accepted that personality variation may affect, and be affected by, an individual’s state. For example, individuals more willing to accept risk (bolder individuals) can have greater access to food (McDonald et al., 2016) and become satiated, while satiation can reduce the risk-taking behavior of bold individuals (Nakayama et al., 2012). These effects can result in feedbacks (Sih et al., 2015) that can be positive (the effect of a behavior on state and the effect of state on behavior act to reinforce each other) or negative (the effect of behavior on state and the effect of state on behavior have opposing effects) (Luttbeg and Sih, 2010; Rands et al., 2003).

Positive feedback can reinforce and magnify personality expression while negative feedback can reduce it. In the context of foraging, negative feedback between state and behavior may occur when individuals with low energy reserves show greater risk-taking behavior than those with high energy reserves and thus acquire more food allowing them to be more cautious in the future (i.e. the asset protection principle (Clark, 1994), although see Rands et al. 2003). However, under high predation risk, positive feedback between state and risk-taking may occur because individuals in good condition are better equipped to escape when confronted with a predator and are therefore willing to take more risks during foraging (i.e. state-dependent safety, Luttbeg and Sih, 2010). In this scenario, individuals in good condition will acquire more resources, reinforcing their condition and helping to maintain personality differences. Despite strong theoretical support, empirical evidence for the effects of state-behavior feedbacks on animal personality is mixed, and mostly confined to studies demonstrating a correlation between state variables and personality traits (Niemelä and Dingemanse, 2018). One exception is a recent study of consistent inter-individual differences in foraging behavior and gizzard mass in red knots, where diet quality was found to increase gizzard mass and larger gizzard size was associated with higher food intake, supporting a positive feedback between gizzard mass and foraging behavior (Mathot et al., 2017).

Fish express consistent inter-individual differences in a range of behaviors related to functionally important tasks, including mating behavior (Magellan and Magurran, 2007), parental care (Budaev et al., 1999), predator avoidance (Kortet et al., 2015) and foraging (MacGregor et al., 2020). Research on the factors that influence the expression of personality in these contexts has mainly focused on the role of predation risk (Brown et al., 2007; Dingemanse et al., 2009; Harris et al., 2010), although increasingly the effects of other environmental variables including abiotic factors are being explored (e.g. temperature (Biro et al., 2010), turbidity (Ehlman et al., 2019), salinity (Sommer-Trembo et al., 2017)), as well as the effects of the social environment (Bevan et al., 2018; McDonald et al., 2016). Together, these studies provide evidence that the expression of consistent individual differences in behavior is highly context dependent, often varying in response to changes in subtle aspects of the environment and over short time scales.

In natural environments food resources fluctuate in space and time; prey species must make decisions whether or not to leave the safety of a refuge to forage (Sih, 1997). These decisions can have enormous impacts on ecological communities owing to their effects on predation risk, predator-prey dynamics, and trophic interactions (Belgrad and Griffen, 2016; Orrock et al., 2013; Sih et al., 1988). In three-spined sticklebacks (*Gasterosteus aculeatus*), refuge use behavior is known to vary consistently between individuals (Bevan et al., 2018; Szopa-Comley et al., 2020) and is a measure of an individual’s willingness to accept potential risk traded-off for greater access to resources (also known as boldness, Balaban-Feld et al., 2019; Harcourt et al., 2009; McDonald et al., 2016). In this study, we presented three-spined sticklebacks with either a foraging context (feeding treatment) or a control trial with no food (control treatment) on alternate days for four consecutive days to experimentally test whether the opportunity to forage affected inter-individual consistency in refuge use behavior. We measured consistency within each pair of repeated time segments: the first five minutes of the two foraging trials, the final five minutes of the foraging trials, the first five minutes of the two control trials, and the final five minutes of the control trials. To examine whether plasticity in refuge use behavior varied with personality, we then tested whether changes in refuge use at the start compared to at the end of the feeding treatment trials differed between bold and shy individuals (as measured in control trials). Actively foraging and consuming food may increase refuge use due to satiation (a negative feedback with boldness) or decrease refuge use as individuals acclimatize more quickly to the area outside the refuge initially perceived as risky (a positive feedback with boldness). If feedback effects are negative due to satiation, we predicted that the opportunity to forage in our feeding treatment would reduce inter-individual consistency in refuge use behavior at the end compared to the start of the trials. In this scenario, bolder individuals were predicted to increase their refuge use because bolder individuals should consume more food and reduce their risk-taking behavior more. If feedback effects are positive due to learning and acclimatization, we predicted that the opportunity to forage would reinforce inter-individual consistency in refuge use behavior, because bolder individuals will learn that their environment offers high reward and low risk, increasing their time spent away from the refuge area. We predicted that feedback effects would be strongest in bolder compared to shyer individuals because bolder individuals will interact more with their environment. Despite the likely role of state-behavior feedbacks in animal personality, evidence for the effects of environmentally induced changes in state on the expression of animal personality is limited. To help address this gap, our study aimed to explicitly test the effect of foraging on the expression of personality differences in refuge use.

## MATERIALS AND METHODS

### Study Animals

Three-spined sticklebacks (37 ± 7.0 mm, standard body length (SL) ± SD at time of testing), were collected from the River Cary, Somerset, UK (ST 469 303) and transported to laboratory facilities. The fish were held for 14 months prior to the experiment in glass tanks (70 cm (L) × 45 cm (W) × 37.5 cm (H)) of approximately 50 individuals each and fed daily with defrosted frozen bloodworm (*Chironomid* larvae). The fish were not sexed because the ambient temperature (16°C) and photocycle (11:13 h light:dark) prevented them from attaining sexual maturation. Sixty-four fish were used in the study.

### Experimental Set-up

Experiments took place in a white acrylic plastic arena (136 (L) × 72 (W) × 19.5 (H) cm) divided into four identical channels (136 (L) × 14.5 (W) cm). Lighting was provided by a florescent lamp positioned at each of the narrow ends of the arena. The arena was sloped lengthways and filled with decholoroniated water varying from 7 cm to 10 cm in depth. In the shallow end of each channel was a single refuge consisting of half a teracotta clay plant pot (10 (L) × 11-7 (W) × 5-3.5 (H) cm) laid on its side. The exit for each refuge faced towards the shallow end wall of the arena and was 15cm from the wall. In the deep end of each channel was a clear petri dish (ø: 9cm) centred 10 cm from the wall so that any food within the petri dish could be visible to the fish once they had exited and swum around the refuge. We filmed trials from above with a GoPro Hero5 video camera (resolution: 1920 × 1080, 30 frames per second) positioned centrally 92 cm above the arena. The camera was connected to an external monitor, allowing observations during trials, and video recording was triggered remotely. The arena was enclosed to camera height with white corregated plastic to minimize external disturbances.

### Experimental protocol

Experiments were conducted on four batches of sixteen fish over four consecutive weeks (23^rd^ October to 16^th^ November 2018). For each fish, testing took place over four consecutive days (Tuesday to Friday). On the Monday morning before the first day of experiments, we assigned sixteen fish to four groups of four individuals and transferred them to smaller glass holding tanks (70 (L) × 25 (W) × 37.5 (H) cm). Assignment was carried out by netting four fish of similar body length from the stock tanks and randomly allocating them to one of the four groups. We repeated this process three more times with different size classes of individuals to create variance in body length within each group that could be used for individual identification. Following assignment, the groups were fed defrosted frozen bloodworm in the afternoon of the same day. Over the subsequent four days, we tested fish once per day with one of two treatment types: feeding or control. All four fish within a group were tested simulutaneously, one fish in each of the four channels. Each batch of 16 fish was alternated as to whether they received the feeding or control treatment on the first day of testing and the order of treatments was then alternated between days. The order of testing of the groups was allocated at random each day within the constraint that each group was tested, first, second, third and forth in their batch over the four days. Each individual in the group was allocated to a channel in the arena at random within the constraint that they experienced all four channels over the four days.

Trials lasted for 40 minutes. In the feeding treatment, fifty medium sized (~1 cm long) bloodworm were placed in the petri dish immediately prior to commencing the trial such that the presense of food could be detected by the fish based on chemical cues but would not be visible until they had exited and swum around the refuge. To quantify food consumption in the feeding treatment, we subtracted the number of bloodworm remaining at the end of the trial from fifty. At the end of a trial the fish were immediately transferred to their holding tank. All groups were fed with bloodworm following the last trial in a day to standardize levels of satiation. The arena water was airated with airstones when not in use. All individuals received four trials except in two cases where two individuals from the same group escaped from their holding tank prior to their second feeding treatment trial. This resulted in a final dataset of 254 trials for 64 individuals. All procedures regarding the use of animals in research followed United Kingdom guidelines and were approved by the institutional ethics committee (UIN UB/17/060).

### Video analysis

Behavioral data were extracted from the video footage using the event recording software BORIS (Friard et al., 2016) by two observers who were allocated trials to process at random and in a random order, and who were blind to the idenities of individual fish during the data extraction. The channels were subdivided along their long axis into three zones: a refuge area, ending at the closed end of the refuge; a neutral area, beginning at the closed end of the refuge and ending on a tangent with the inner edge of the petri dish; and a feeding area, beginning on a tangent with the inner end of the petri dish and ending at the wall. We quantified the following behaviors from the videos: latency to emerge from the refuge that terminated once the fish had their entire body out of the refuge, which we used to measure boldness (e.g. Brown et al., 2005); the duration of time (to the nearest second) that the fish spent in the refuge area for the start and end five minute segments of the trial, where we deemed that a fish had crossed from one zone to another when their head crossed the boundary between zones; and whether a fish fed in each of the start and end five minute segments of a trial. If the fish did not emerge during the trial they were given an emergence latency of 2400 s to match the length of the trial. One fish was not successfully transferred into the refuge at the start of the second feeding treatment trial and was therefore excluded from analyses of inter-individual consistency in latency to emerge from the refuge.

### Statistical analyses

Statistical analyses were performed in R version 3.6.0. The initial analyses tested whether the willingness to accept risk and emerge from the refuge was affected by experimental variables (treatment and trial number) and body length. A generalized linear mixed model (GLMM) with binomial error distribution was used to test for the effects of treatment, trial number (1 to 4), and standard body length (SL) on the likelihood that a fish emerged from the refuge during a trial (coded 0: no emergence or 1: emerged) with individual identity included as a random intercept. A negative binomial GLMM including treatment, trial number, and SL as main effects and individual identity as a random intercept was used to examine the predictors of latency to emerge from the refuge. To test whether individual identity accounted for significant variation in the likelihood and the latency of fish to emerge from the refuge, we compared the goodness-of-fit (deviance) of the GLMMs to the models with individual identity removed using a likelihood ratio test (LRT).

To estimate inter-individual consistency in the latency to emerge from the refuge (with a maximum value of 2400 s) in the control and feeding treatments and the time spent in the refuge area during the start and end five minute periods (with a maximum value of 300 s assigned for each time segment) we used Spearman’s rank correlation coefficients due to the statistical issues associated with a large proportion of data points being right-censored (e.g. inflated repeatability, Stamps et al., 2012). To statistically compare the correlation coefficients we performed randomization tests with 1,000 iterations (Manly, 1991). For emergence latency, the Spearman’s rank correlation coefficient was calculated separately for the control and feeding treatment trials. The difference between these correlations was used as the observed difference in inter-individual consistency in emergence latency between treatments. For each iteration of the randomization, each individual fish’s emergence latencies were randomly shuffled between treatments, and the correlation coefficients, and their difference, was recalculated. We compared the absolute observed difference of the correlation coefficients to the frequency distribution of the absolute randomized expected differences to determine the significance (alpha = 0.05). We used a similar approach to compare the Spearman’s rank correlation coefficient in time spent in the refuge area between the start segments of the two feeding trials, and separately the two end segments. Here, the difference between the start and end correlation coefficients was used as the observed change in consistency between the start and end segments. For each iteration of the randomization, each individual fish’s values for the time spent in the refuge area were randomly shuffled between the start and the end segments, and the correlation coefficients, and their difference, was recalculated. The analysis was repeated for the control treatment trials. The feeding trial data were then split between the 50% boldest and 50% shyest fish, and the randomisation procedure repeated on each subset of the data separately. Individuals were catagorised as bold (n = 31) or shy (n = 31) based on their mean emergence latency from the refuge in the two control treatment trials with the median value (246.75 s) across individuals as the cut-off threshold between the categories (n = 2 individuals were excluded due to missing data).

An individual’s mean latency to emerge from the refuge in the control treatment trials was used as an estimate of their boldness in further analyses (where smaller values represent bolder fish). To test whether the absolute change or directional change in individuals’ refuge use during the feeding trials was associated with boldness, we estimated the Spearman’s rank correlation between the mean latency to emerge from the refuge in the control treatment trials and the absolute (i.e. negative values made positive) difference in the time that a fish spent in the refuge area between the start and end of the feeding treatments, using an individual’s mean difference in time from the two trials. The correlation test then was repeated using the non-absolute rather than absolute difference. GLMMs were not performed due to violation of parametric assumptions.

To test whether boldness predicted foraging behavior we used Generalized linear models (GLM) with binomial error distribution to test whether the mean latency to emerge from the refuge in the control treatment predicted the likelihood that a fish fed during the start, and in a separate model at the end, of the feeding treatment, controlling for SL and test order (1st or 2nd) as main effects. Individual identity was not included in the models because the random effect variance was estimated close to zero. The two individuals with data for only one trial were excluded from both analyses. The likelihood that a fish fed was analyzed as a response variable rather than the number of bloodworm consumed per individual because in over half of the trials no bloodworm were consumed by the focal fish at the start and at the end of the feeding treatment.

Analyses assuming a negative binomial distribution were checked for model assumptions using diagnostic plots in R package *DHARMa*. The statistical significance of fixed effects was tested with likelihood ratio tests in R package *lme4*.

## RESULTS

### Inter-individual consistency in risk-taking behavior

In 13% of cases (n = 33/253) the fish did not emerge from the refuge during a trial. There was no significant difference between the feeding and control treatments (GLMM (binomial): treatment (control as reference level): Estimate = 0.52 ± 0.461, χ^2^ = 1.32, *P* = 0.25), or between fish of different body length (SL: Estimate = −0.062 ± 0.0520, χ^2^ = 1.43, *P* = 0.23), in their likelihood to emerge from the refuge. However, there was a significant effect of trial number with the likelihood that a fish emerged from the refuge declining significantly over the four trials (trial number: Estimate = −0.54 ± 0.218, χ^2^ = 6.79, *P* = 0.009). There were consistent differences between individuals in their likelihood to emerge, including after controlling for body length, treatment, and trial number (Individual Identity Intercept: LRT: χ^2^ = 15.6, *P* < 0.0001).

The latency to first leave the refuge during a trial was longer for larger fish than for smaller fish (GLMM (negative binomial): SL: Estimate = 0.06 ± 0.021, χ^2^ = 7.90, *P* = 0.05) and increased over the four days of trials (trial number: Estimate = 0.23 ± 0.059, χ^2^ = 15.28, *P* < 0.0001), but there was no significant effect of treatment (treatment (control as reference level): Estimate = 0.07 ± 0.119, χ^2^ = 0.39, *P* = 0.53). There were consistent differences between individuals in their latency to first leave the refuge, including after controlling for body length, treatment, and trial number (Individual Identity Intercept: LRT: χ^2^ = 89.4, *P* < 0.0001).

When analyzing the data from the two treatments separately, inter-individual differences in latency to emerge from the refuge were significantly correlated in the control (Spearman’s rank correlation: R_s_ = 0.53, p < 0.0001, n = 61) and feeding (Spearman’s rank correlation: R_s_ = 0.62, p < 0.0001, n = 61) treatments, however the difference between treatments in the Spearman’s rank correlation coefficients in latency to emerge from the refuge did not differ significantly from the frequency distribution of the absolute randomized expected differences suggesting there was no significant effect of treatment on the consistency in inter-individual differences (observed absolute difference in Spearman’s rank correlation = 0.10, mean expected absolute difference in Spearman’s rank correlation = 0.10, *P* = 0.50, Fig. S1). An individual’s mean latency to emerge from the refuge in the control treatment (boldness) was significantly positively correlated with their mean latency to emerge from the refuge in the feeding treatment (Spearman’s rank correlation: R_s_ = 0.75, *P* < 0.0001, n = 64).

### Effects of opportunity to forage and personality type on inter-individual consistency in refuge use

Inter-individual differences in the time spent in the refuge area were significantly correlated in the start (Spearman’s rank correlation: R_s_ = 0.73, p < 0.0001, n = 62) and end (R_s_ = 0.43, *P* = 0.0006, n = 62) segments of the two feeding treatment trials and almost significantly correlated at the start (R_s_ = 0.24, *P* = 0.06, n = 64) and significantly correlated at the end (R_s_ = 0.33, *P* = 0.008, n = 64) segments of the two control treatment trials (Fig. 1). The difference in the Spearman’s rank correlation coefficients in the time spent in the refuge area across the start of the trials and across the end of the trials did not differ significantly from the frequency distribution of the absolute randomized expected differences in the control treatment (observed absolute difference in Spearman’s rank correlation = 0.09, mean expected absolute difference in Spearman’s rank correlation = 0.13, *P* = 0.58, Fig. 2a). In contrast, in the feeding treatment the difference between the correlation coefficients did differ significantly from expected (observed absolute difference in Spearman’s rank correlation = 0.30, mean expected absolute difference in Spearman’s rank correlation = 0.11, *P* = 0.021, Fig. 2b). The time in the refuge area was less correlated between the two feeding trials at the end compared to the start of the trials, i.e. the correlation decreased between the start and end segments (Fig. 1b, d).

**Figure 1.**
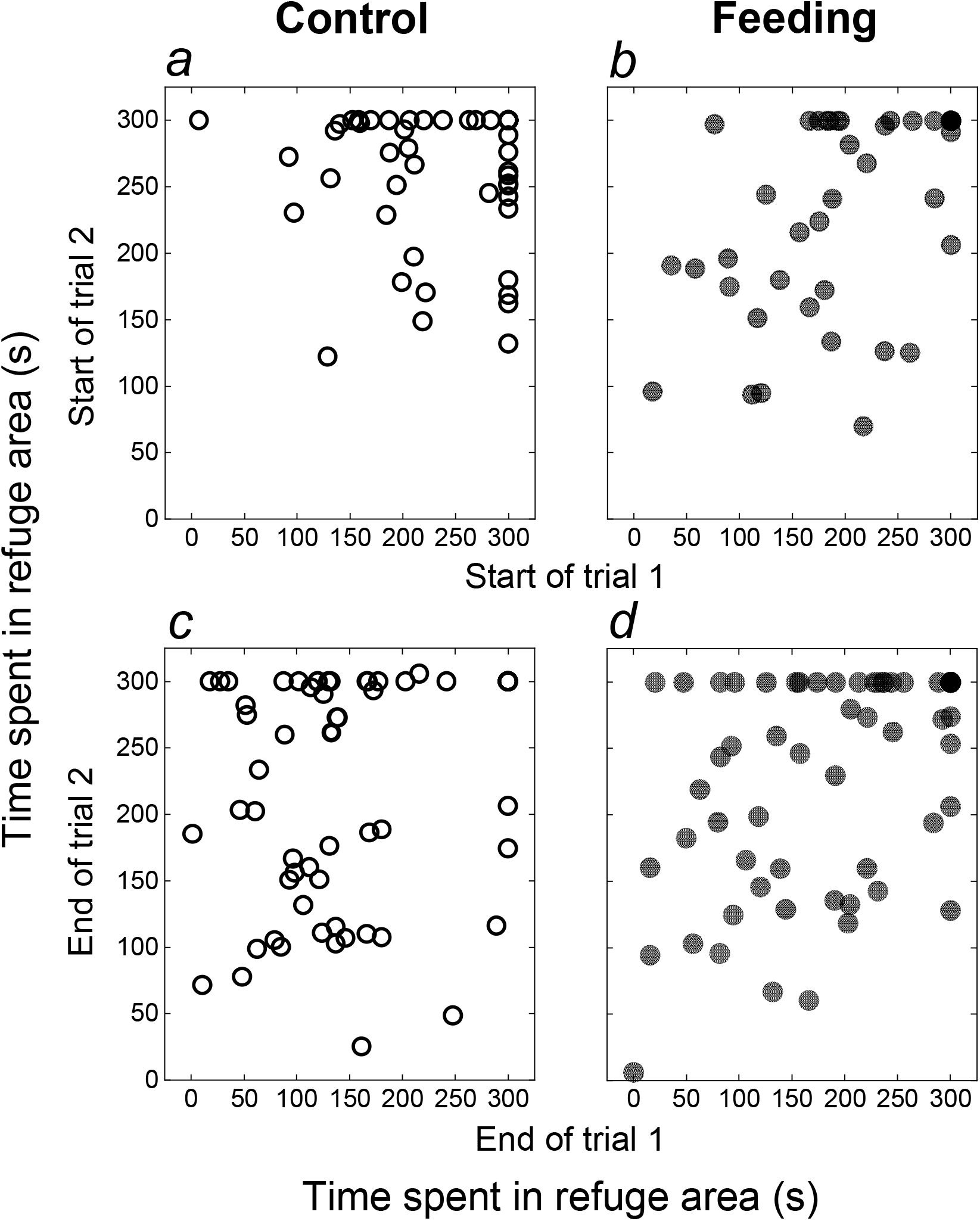
Correlations between the time spent in the refuge area at the start (a, b) and at the end (c, d) of the trials for the control (a, c) and feeding (b, d) treatments. Points depict data for individual fish (white: control treatment and black: feeding treatment).

**Figure 2.**
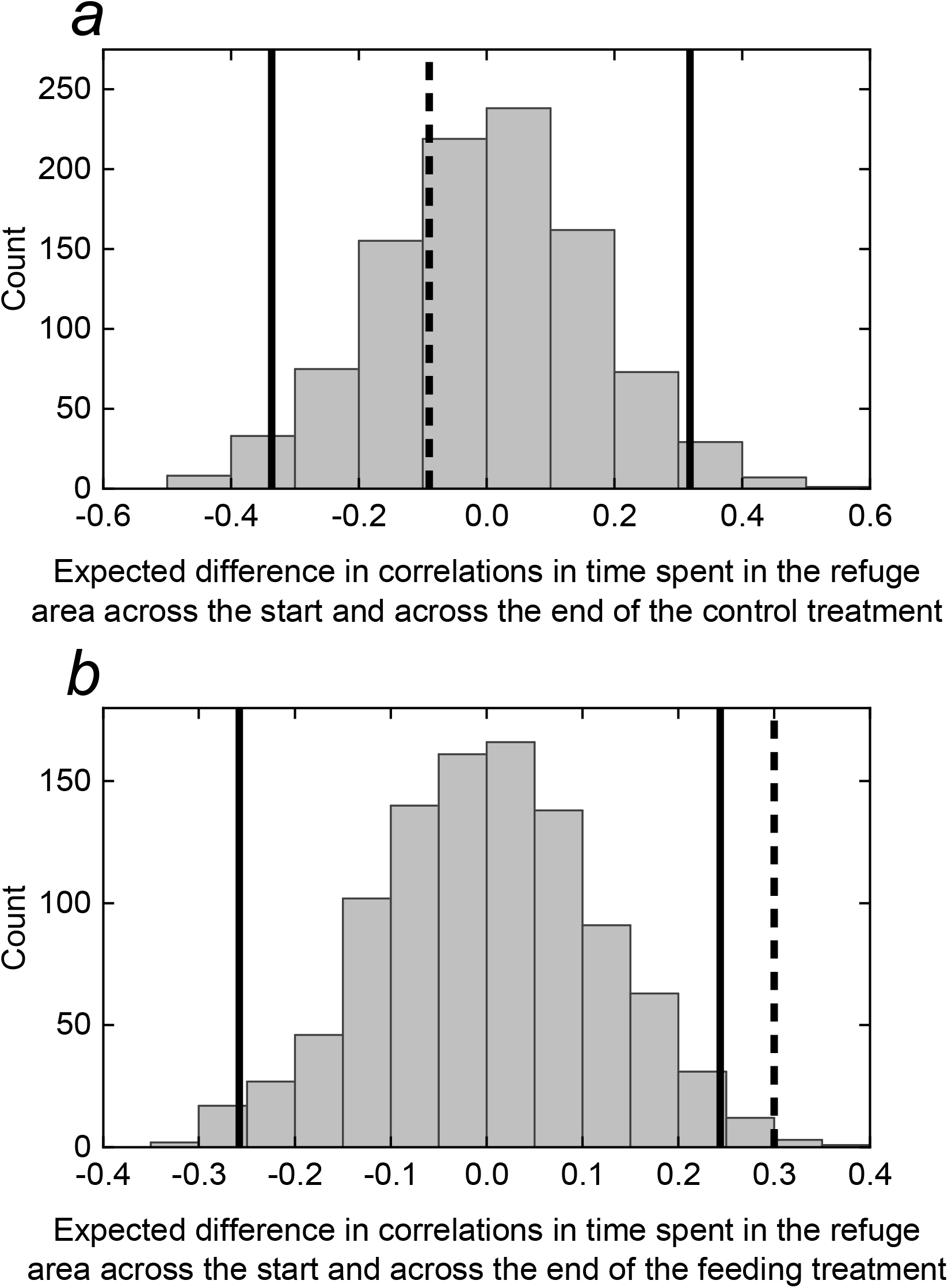
The expected difference in Spearman’s rank correlation coefficients in the time spent in the refuge area across the start and across the end of the trials for (a) the control treatment and (b) the feeding treatment. Values are based on 1,000 randomizations of the data within individuals. If the observed difference (dashed black line) is outside of the 95% limits of the randomized data’s distribution (solid black lines), it is unlikely that observed difference occurred by chance.

The decreased inter-individual consistency in refuge use at the end of the feeding treatment trials appeared to be driven by the behavior of the bold fish, which were more consistent in their refuge use at the start (Spearman’s rank correlation: R_s_ = 0.70, p < 0.001, n = 31) than at the end (R_s_ = 0.37, *P* = 0.038, n = 31) of the feeding trials compared to the shy fish, which were less consistent at the start (R_s_ = 0.30, *P* = 0.10, n = 31) than at the end (R_s_ = 0.46, *P* = 0.01, n = 31; Fig. S2). However, the difference in correlations for the start and end segments were not significantly different from expected in the bold fish (Feeding Treatment: Observed absolute difference in bold fish: R_s_ = 0.33, mean expected absolute difference in R_s_ in bold fish: R_s_ = 0.16, *P* = 0.096) or in the shy fish (Observed difference in shy fish: R_s_ = 0.15, mean expected difference in R_s_ in shy: R_ind_ = 0.20, *P* = 0.54, no. of randomizations = 1,000, Fig. S3).

During the feeding treatment trials, the mean absolute difference in time that fish spent in the refuge area between the start and end of the feeding treatments was significantly negatively correlated with their mean latency to emerge from the refuge in the control treatment (Spearman’s rank correlation: R_s_ = −0.40, *P* = 0.0012, n = 62, Fig. S4). Bold fish changed their refuge use behavior more between the start and end of the feeding treatment trials than shy fish (Fig. 3). However, there was no significant correlation between the mean latency to emerge from the refuge in the control treatment and the mean non-absolute difference in time spent in the refuge area between the start and end of the feeding treatments (Spearman’s rank correlation: R_s_ = 0.18, *P* = 0.16, n = 62). While shy fish tended to not change their refuge use at the end compared to the start of the feeding treatment trials, bold fish showed more variation with some bold individuals increasing and others decreasing their refuge use at the end compared to the start of the trials (Fig. 4).

**Figure 3.**
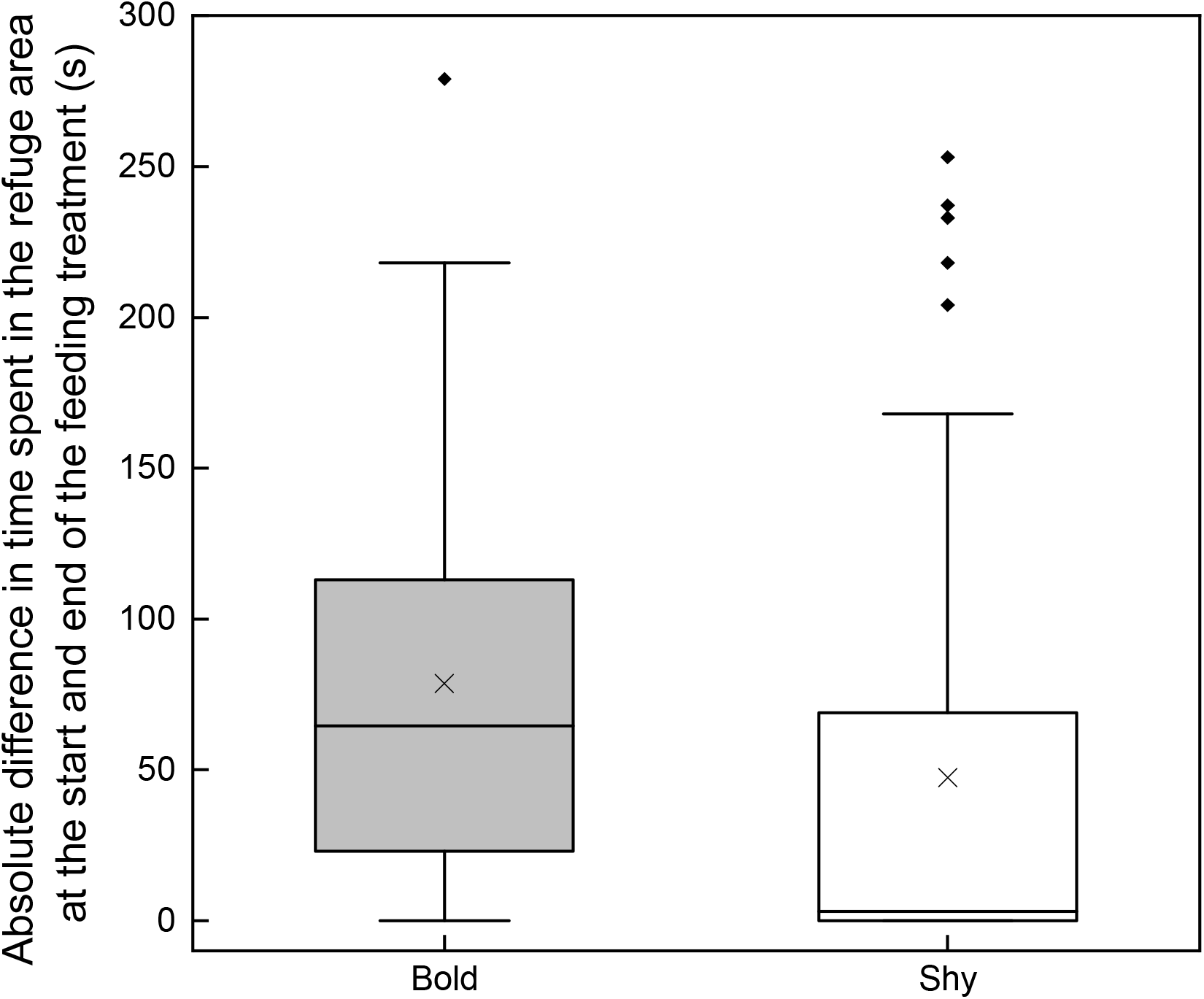
The absolute difference in the time spent in the refuge area at the start and end of the feeding treatment for bold and shy fish. The boxes depict the interquartile range of the data and the median, and the whiskers extend to 1.5 × the interquartile range. Data beyond the whiskers are shown as points. The crosses indicate the mean value. Fish were categorized as bold or shy based on their mean latency to emerge from the refuge in control treatment (threshold set as the median value: 246.75 s).

**Figure 4.**
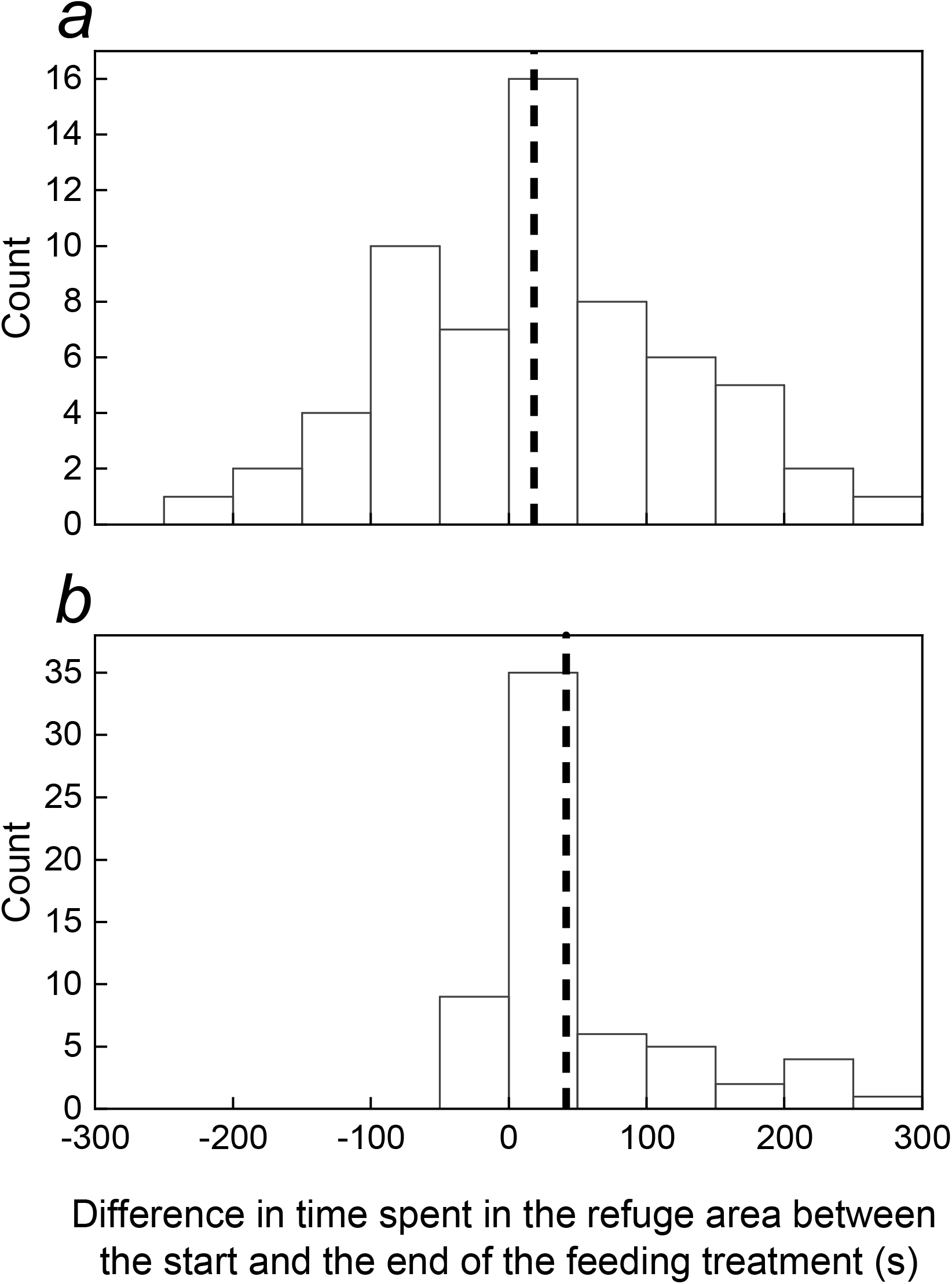
Difference in time spent in the refuge area between the start and end of the feeding treatment in (a) bold fish and (b) shy fish. The black dashed line shows the mean difference in time in each case. Fish were categorized as bold or shy based on their mean latency to emerge from the refuge in control treatment (threshold set as the median value: 246.75 s).

### Predictors of bloodworm consumption

Individuals consumed 17 ± 1.2 (mean ± standard error) bloodworm on average in the feeding treatment trials (Fig. S5). In only 4 of 126 trials were all 50 bloodworm consumed, and in 23 trials no bloodworm were consumed. Changes in refuge use between the start and the end of a trial may be driven by food consumed, either due to positive feedback (e.g. learning about the food reward outside of the refuge) or negative feedback (e.g. from satiation). The likelihood that a fish fed during the start of a feeding trial was predicted by boldness (GLM (binomial): mean latency to emerge from refuge in control treatment: Estimate = −0.95 ± 0.337, χ^2^ = 11.69, *P* = 0.0006), but indicative of satiation, this was not the case at the end of the feeding trials (mean latency to emerge from refuge in control treatment: Estimate = 0.15 ± 0.280, χ^2^ = 0.29, *P* = 0.589). There was also a significant effect of body size on the likelihood of feeding at the start with smaller fish more likely to feed (SL: Estimate = −0.58 ± 0.222, χ^2^ = 7.40, *P* = 0.0064), however, there was no effect of body size at the end of the trials (SL: Estimate = −0.43 ± 0.307, χ^2^ = 2.12, *P* = 0.15). Test order was not a significant predictor of the likelihood that a fish fed at either the start (test order (1st test as reference level): Estimate = −0.52 ± 0.307, χ^2^ = 1.54, *P* = 0.21) or the end (test order (1st test as reference level): Estimate = 0.29 ± 0.554, χ^2^ = 0.29, *P* = 0.59) of the trials.

## DISCUSSION

Here we show that consistent inter-individual differences in refuge use in three-spined sticklebacks were reduced following the opportunity to forage, suggesting that foraging can suppress the expression of personality variation. Although bold fish were less correlated in their refuge use at the end compared to the start of the feeding treatment trials the difference was not statistically significant. However, as in studies of sticklebacks demonstrating that plasticity in behavior can covary with personality (e.g. Bevan et al., 2018; Harcourt et al., 2009; Laskowski and Bell, 2013), we found that bold fish changed their refuge use more than shy fish between the start and end of the feeding treatment trials. This greater change in the behaviour of bolder fish was predicted from state-behaviour feedbacks as bold fish are more likely to interact with the environment. An individual’s boldness predicted whether they fed at the start, but not at the end, of the feeding treatment trials in support of negative feedback between state (satiation) and behavior (refuge use) causing the observed reduction in personality expression. However, contrary to predictions of negative feedbacks, we did not find evidence that bolder fish used the refuge more at the end compared to the start of the trials, with some bold individuals increasing and others decreasing their refuge use.

While previous studies have often sought to understand how individuals’ personality traits change over long time frames, such as during ontogeny (Brommer and Class, 2015), very little is known about the processes underlying short-term fluctuations in personality expression. The state dependency of behavior is a central concept in theory to explain the origin and maintenance of consistent behavioral variation but could also provide a mechanistic explanation for breakdowns in personality expression when state variables are labile and produce a negative feedback on behavior (Dingemanse and Wolf, 2010; Sih et al., 2015). In support of this, reduced inter-individual consistency in refuge use during the feeding trials was associated with the opportunity for individuals to increase their nutritional state, and no change in inter-individual consistency was observed in control trials without food. Furthermore, the absence of a significant difference in the first time to leave the refuge (i.e. the emergence latency) between the treatments suggested that it was encountering the food directly rather than the presence of food olfactory cues that affected the inter-individual consistency in behavior.

Previous studies on fish have shown that bolder individuals are more likely to feed in risky contexts (McDonald et al., 2016) but are also more at risk of predation (Balaban-Feld et al., 2019; Bell and Sih, 2007), and that nutritional state can influence foraging behavior (Salvanes and Hart, 1998). Therefore, we predicted that a satiation effect in the feeding treatment trials would be strongest in bolder individuals, and, as a result, bolder individuals would change their behavior more than shy individuals and in the direction of increased refuge use at the end compared to the start of the trials (due to being satiated and the potential risks associated with being away from the refuge). However, counter to expectation, our results did not support directionality to the behavioral changes in bolder individuals, with some bolder individuals expressing even bolder behavior (more time away from the refuge) and others converging on the refuge use behavior of shy fish (as predicted due to negative feedback effects between satiation and refuge use). Some of this variation among bold fish could be explained by variation in body size, because smaller fish should satiate more quickly (Brett, 1971; Ende et al., 2018). However, while we found that smaller fish were more likely to feed at the start of the trials than larger fish, there was no body size effect at the end of trials. An alternative explanation is that the motivation to feed away from a refuge over repeated foraging trips close together in time initially increases with acclimatization and learning about the resource, then reduces with satiation. Such a trend over time was documented by McDonald et al. (McDonald et al., 2016) in shoals of three-spined sticklebacks. In our experiment, some bold fish may have been in the first phase, reducing their refuge use at the end of the trials, while others were in the satiation phase, increasing their refuge use at the end of the trials.

Spatial and temporal fluctuations in the availability of food resources are common in natural environments (Ward et al., 2006). Our results show that individual differences between bold fish may become less consistent when resources are abundant, resulting in the suppression of personality variation. This suggests that the maintenance of personality types could depend on foraging opportunities, with consistent individual differences in behavior more strongly expressed in low resource environments. Such inconsistency in behavior during foraging could also have an important adaptive function, for example, by making individuals less predictable to predators or competitors if encountering the same individuals repeatably (Briffa, 2013; Chang et al., 2017), weakening the strength of directional selection on boldness behavior and its correlated traits. More generally, inconsistency in the expression of phenotypes may have important evolutionary consequences by potentially weakening evolutionary responses to changes in the environment. Future work that considers the context-dependent expression of animal personality will help to better understand the selection pressures that shape consistent inter-individual differences in behavior. There is growing evidence of how boldness behavior such as refuge use can have ecological consequences, including influences on population dynamics via individuals’ growth, survival, and reproductive success, and on trophic interactions via the effects on the costs and benefits of different predator strategies (Belgrad and Griffen, 2016; Orrock et al., 2013; Sih et al., 1988). However, one key outstanding question is in the ecological implications of covariation between personality and behavioral plasticity. Our study suggests this could be particularly challenging to address when behavioral plasticity is unpredictable due to unpredictable state-behavior feedbacks.

## Acknowledgements

This work was supported by the Natural Environment Research Council grant number NE/P012639/1 to C.C.I. We also thank Andrew Szopa-Comley for comments and Sarah Washington for help carrying out the experiment.

## REFERENCES

Balaban-Feld J, Mitchell WA, Kotler BP, Vijayan S, Elem LTT, Rosenzweig ML, Abramsky Z, 2019. Individual willingness to leave a safe refuge and the trade-off between food and safety: a test with social fish. Proceedings of the Royal Society B: Biological Sciences 286:20190826.

Belgrad BA, Griffen BD, 2016. Predator and prey interactions mediated by prey personality and predator hunting mode. Proceedings of the Royal Society B: Biological Sciences 283:20160408.

Bell AM, Sih A, 2007. Exposure to predation generates personality in threespined sticklebacks (Gasterosteus aculeatus). Ecology Letters 10:828–834.

Bevan PA, Gosetto I, Jenkins ER, Barnes I, Ioannou CC, 2018. Regulation between personality traits: individual social tendencies modulate whether boldness and leadership are correlated. Proceedings of the Royal Society B: Biological Sciences 285:20180829.

Biro PA, Beckmann C, Stamps JA, 2010. Small within-day increases in temperature affects boldness and alters personality in coral reef fish. Proceedings of the Royal Society B: Biological Sciences 277:71–77.

Boissy A, 1995. Fear and Fearfulness in Animals. The Quarterly Review of Biology 70:165–191.

Boon AK, Reale D, Boutin S, 2007. The interaction between personality, offspring fitness and food abundance in North American red squirrels. Ecology Letters 10:1094–1104.

Brett JR, 1971. Satiation Time, Appetite, and Maximum Food Intake of Sockeye Salmon (Oncorhynchus nerka). Journal of the Fisheries Research Board of Canada 28:409–415.

Briffa M, 2013. Plastic proteans: reduced predictability in the face of predation risk in hermit crabs. Biol Lett 9:20130592–20130592.

Briffa M, Rundle SD, Fryer A, 2008. Comparing the strength of behavioural plasticity and consistency across situations: animal personalities in the hermit crab Pagurus bernhardus. Proceedings of the Royal Society B: Biological Sciences 275:1305–1311.

Brommer JE, Class B, 2015. The importance of genotype-by-age interactions for the development of repeatable behavior and correlated behaviors over lifetime. Frontiers in Zoology 12:S2.

Brown C, Burgess F, Braithwaite VA, 2007. Heritable and experiential effects on boldness in a tropical poeciliid. Behavioral Ecology and Sociobiology 62:237–243.

Brown C, Jones F, Braithwaite V, 2005. In situ examination of boldness–shyness traits in the tropical poeciliid, Brachyraphis episcopi. Animal Behaviour 70:1003–1009.

Budaev SV, Zworykin DD, Mochek AD, 1999. Individual differences in parental care and behaviour profile in the convict cichlid: a correlation study. Animal Behaviour 58:195–202.

Carere C, Gherardi F, 2013. Animal personalities matter for biological invasions. Trends in Ecology & Evolution 28:5–6.

Chang C-c, Teo HY, Norma-Rashid Y, Li D, 2017. Predator personality and prey behavioural predictability jointly determine foraging performance. Scientific Reports 7:40734.

Clark CW, 1994. Antipredator behavior and the asset-protection principle. Behavioral Ecology 5:159–170.

Dall SRX, Bell AM, Bolnick DI, Ratnieks FLW, 2012. An evolutionary ecology of individual differences. Ecology Letters 15:1189–1198.

Dall SRX, Houston AI, McNamara JM, 2004. The behavioural ecology of personality: consistent individual differences from an adaptive perspective. Ecology Letters 7:734–739.

Dingemanse JN, Réale D, 2005. Natural selection and animal personality. Behaviour 142:1159–1184.

Dingemanse NJ, Bouwman KM, van de Pol M, van Overveld T, Patrick SC, Matthysen E, Quinn JL, 2012. Variation in personality and behavioural plasticity across four populations of the great tit Parus major. Journal of Animal Ecology 81:116–126.

Dingemanse NJ, Kazem AJ, Reale D, Wright J, 2010. Behavioural reaction norms: animal personality meets individual plasticity. Trends in Ecology & Evolution 25:81–89.

Dingemanse NJ, Van der Plas F, Wright J, Reale D, Schrama M, Roff DA, Van der Zee E, Barber I, 2009. Individual experience and evolutionary history of predation affect expression of heritable variation in fish personality and morphology. Proceedings of the Royal Society B: Biological Sciences 276:1285–1293.

Dingemanse NJ, Wolf M, 2010. Recent models for adaptive personality differences: a review. Philosophical Transactions of the Royal Society B: Biological Sciences 365:3947–3958.

Ehlman SM, Halpin R, Jones C, Munson A, Pollack L, Sih A, 2019. Intermediate turbidity elicits the greatest antipredator response and generates repeatable behaviour in mosquitofish. Animal Behaviour 158:101–108.

Ende SSW, Thiele R, Schrama JW, Verreth JAJ, 2018. The influence of prey density and fish size on prey consumption in common sole (Solea solea L.). Aquatic Living Resources 31:16.

Friard O, Gamba M, Fitzjohn R, 2016. BORIS: a free, versatile open-source event-logging software for video/audio coding and live observations. Methods in Ecology and Evolution 7:1325–1330.

Harcourt JL, Ang TZ, Sweetman G, Johnstone RA, Manica A, 2009. Social feedback and the emergence of leaders and followers. Current Biology 19:248–252.

Harris S, Ramnarine IW, Smith HG, Pettersson LB, 2010. Picking personalities apart: estimating the influence of predation, sex and body size on boldness in the guppy Poecilia reticulata. Oikos 119:1711–1718.

Houston AI, McNamara JM, I HA, 1999. Models of Adaptive Behaviour: An Approach Based on State: Cambridge University Press.

Huntingford FA, 1976. The relationship between anti-predator behaviour and aggression among conspecifics in the three-spined stickleback, Gasterosteus Aculeatus. Animal Behaviour 24:245–260.

Kortet R, Sirkka I, Lai Y-T, Vainikka A, Kekäläinen J, 2015. Personality differences in two minnow populations that differ in their parasitism and predation risk. Frontiers in Ecology and Evolution 3:1–9.

Laskowski KL, Bell AM, 2013. Competition avoidance drives individual differences in response to a changing food resource in sticklebacks. Ecology Letters 16:746–753.

Luttbeg B, Sih A, 2010. Risk, resources and state-dependent adaptive behavioural syndromes. Philosophical Transactions of the Royal Society B: Biological Sciences 365:3977–3990.

MacGregor HEA, Herbert-Read JE, Ioannou CC, 2020. Information can explain the dynamics of group order in animal collective behaviour. Nature Communications 11:2737.

Magellan K, Magurran AE, 2007. Behavioural profiles: individual consistency in male mating behaviour under varying sex ratios. Animal Behaviour 74:1545–1550.

Magurran AE, May RM, Wilson DS, 1998. Adaptive individual differences within single populations. Philosophical Transactions of the Royal Society of London Series B: Biological Sciences 353:199–205.

Mangel M, 1991. Adaptive walks on behavioural landscapes and the evolution of optimal behaviour by natural selection. Evolutionary Ecology 5:30–39.

Manly BFJ, 1991. Randomization and Monte Carlo methods in biology: Chapman and Hall.

Mathot KJ, Dekinga A, Piersma T, Sandercock B, 2017. An experimental test of state–behaviour feedbacks: gizzard mass and foraging behaviour in red knots. Functional Ecology 31:1111–1121.

Mathot KJ, van den Hout PJ, Piersma T, Kempenaers B, Reale D, Dingemanse NJ, 2011. Disentangling the roles of frequency-vs. state-dependence in generating individual differences in behavioural plasticity. Ecology Letters 14:1254–1262.

McDonald ND, Rands SA, Hill F, Elder C, Ioannou CC, 2016. Consensus and experience trump leadership, suppressing individual personality during social foraging. Science Advances 2:e1600892.

Nakayama S, Johnstone RA, Manica A, 2012. Temperament and Hunger Interact to Determine the Emergence of Leaders in Pairs of Foraging Fish. PLOS ONE 7:e43747.

Niemelä PT, Dingemanse NJ, 2018. Meta-analysis reveals weak associations between intrinsic state and personality. Proceedings of the Royal Society B: Biological Sciences 285:20172823.

Oers Kv, Drent PJ, Goede Pd, Noordwijk AJv, 2004. Realized heritability and repeatability of risk-taking behaviour in relation to avian personalities. Proceedings of the Royal Society of London Series B: Biological Sciences 271:65–73.

Orrock JL, Preisser EL, Grabowski JH, Trussell GC, 2013. The cost of safety: Refuges increase the impact of predation risk in aquatic systems. Ecology 94:573–579.

Rands AS, Cowlishaw G, Pettifor AR, Rowcliffe JM, Johnstone AR, 2003. Spontaneous emergence of leaders and followers in foraging pairs. Nature 423:432–434.

Salvanes AGV, Hart PJB, 1998. Individual variability in state-dependent feeding behaviour in three-spined sticklebacks. Animal Behaviour 55:1349–1359.

Sih A, 1997. To hide or not to hide? Refuge use in a fluctuating environment. Trends in Ecology & Evolution 12:375–376.

Sih A, Bell A, Johnson JC, 2004. Behavioral syndromes: an ecological and evolutionary overview. Trends in Ecology & Evolution 19:372–378.

Sih A, Mathot KJ, Moiron M, Montiglio PO, Wolf M, Dingemanse NJ, 2015. Animal personality and state-behaviour feedbacks: a review and guide for empiricists. Trends in Ecology & Evolution 30:50–60.

Sih A, Petranka JW, Kats LB, 1988. The dynamics of prey refuge use: A model and tests with sunfish and salamander larvae. The American naturalist 132:463–483.

Smith BR, Blumstein DT, 2008. Fitness consequences of personality: a meta-analysis. Behavioral Ecology 19:448–455.

Snell-Rood EC, 2013. An overview of the evolutionary causes and consequences of behavioural plasticity. Animal Behaviour 85:1004–1011.

Sommer-Trembo C, Petry AC, Gomes Silva G, Vurusic SM, Gismann J, Baier J, Krause S, Iorio JdAC, Riesch R, Plath M, 2017. Predation risk and abiotic habitat parameters affect personality traits in extremophile populations of a neotropical fish (Poecilia vivipara). Ecology and Evolution 7:6570–6581.

Stamps JA, 2016. Individual differences in behavioural plasticities. Biological Reviews 91:534–567.

Stamps JA, Briffa M, Biro PA, 2012. Unpredictable animals: individual differences in intraindividual variability (IIV). Animal Behaviour 83:1325–1334.

Stoffel MA, Nakagawa S, Schielzeth H, Goslee S, 2017. rptR: repeatability estimation and variance decomposition by generalized linear mixed-effects models. Methods in Ecology and Evolution 8:1639–1644.

Szopa-Comley AW, Donald WG, Ioannou CC, 2020. Predator personality and prey detection: inter-individual variation in responses to cryptic and conspicuous prey. Behavioral Ecology and Sociobiology 74:70.

Via S, Gomulkiewicz R, De Jong G, Scheiner SM, Schlichting CD, Van Tienderen PH, 1995. Adaptive phenotypic plasticity: consensus and controversy. Trends in Ecology & Evolution 10:212–217.

Ward AJW, Webster MM, Hart PJB, 2006. Intraspecific food competition in fishes. Fish and Fisheries 7:231–261.

Westneat DF, Hatch MI, Wetzel DP, Ensminger AL, 2011. Individual Variation in Parental Care Reaction Norms: Integration of Personality and Plasticity. The American naturalist 178:652–667.

