## Supplementary figures and tables for "State-behavior feedbacks suppress personality variation in boldness during foraging in sticklebacks"

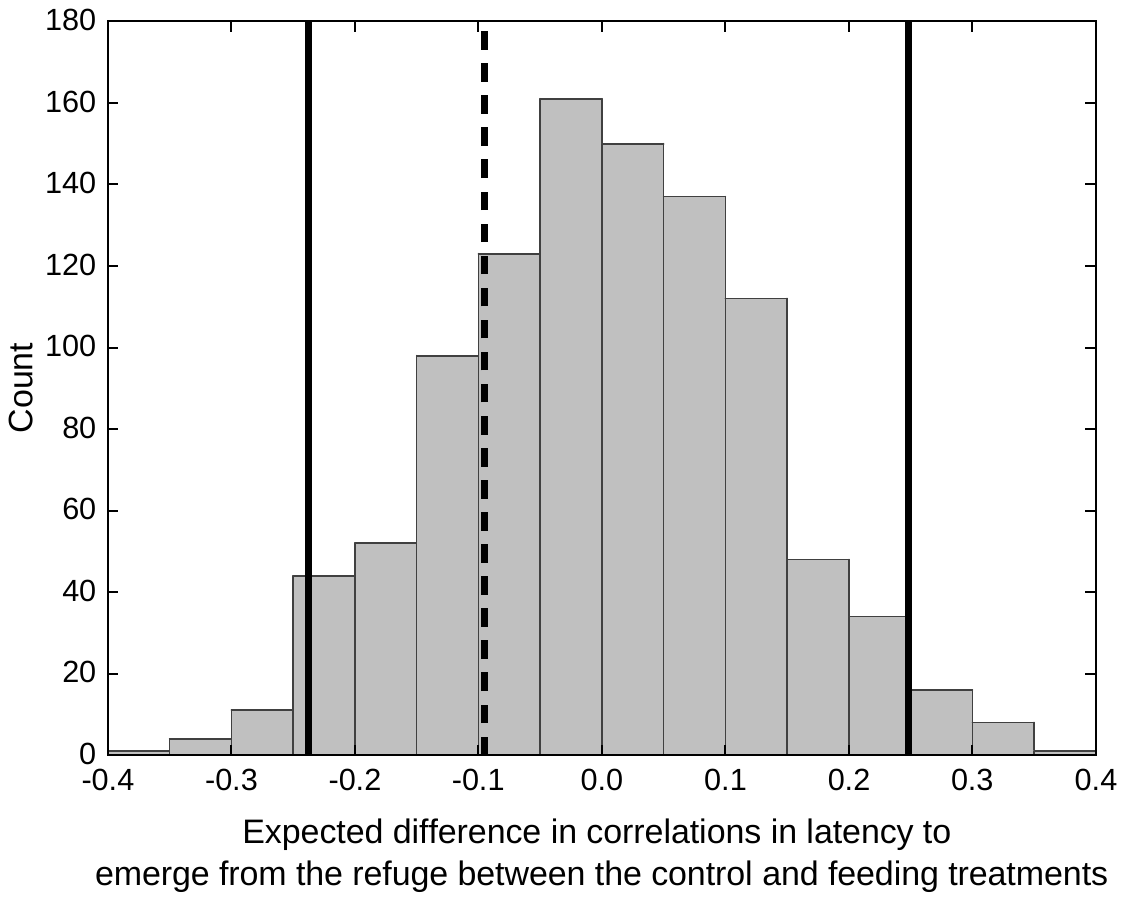

**Figure S1**. The expected difference in Spearman’s rank correlation coefficients in the latency to emerge from the refuge between the control and feeding treatments. Values are based on 1,000 randomizations of the data within individuals. If the observed difference (dashed black line) is outside of the 95% limits of the randomized data’s distribution (solid black lines), it is unlikely that observed difference occurred by chance.

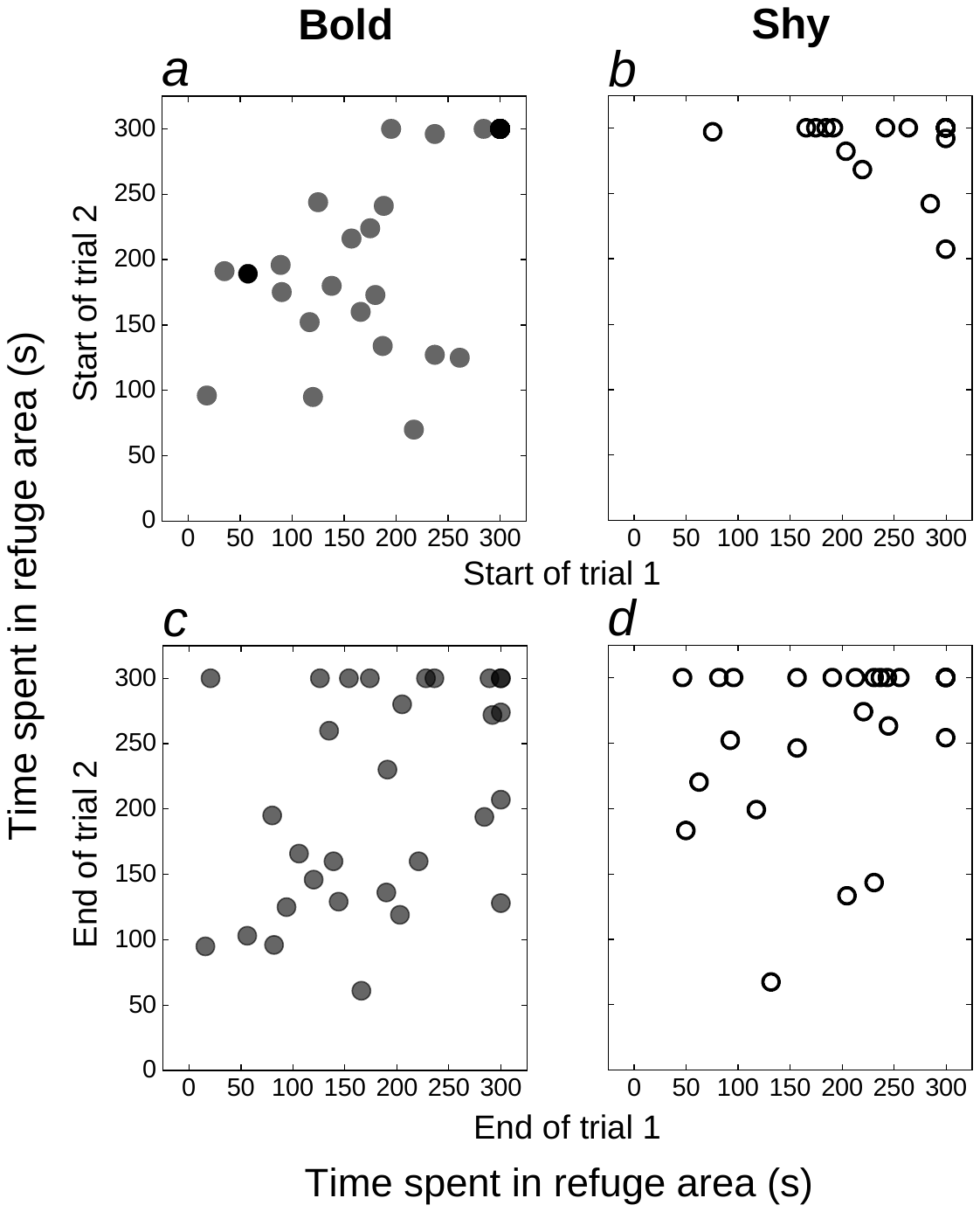

**Figure S2**. Correlations between the time spent in the refuge area at the start (a, b) and at the end (c, d) of the feeding treatment trials for bold (a, c) and shy (b, d) fish. Points depict data for individual fish (white: control treatment and black: feeding treatment).

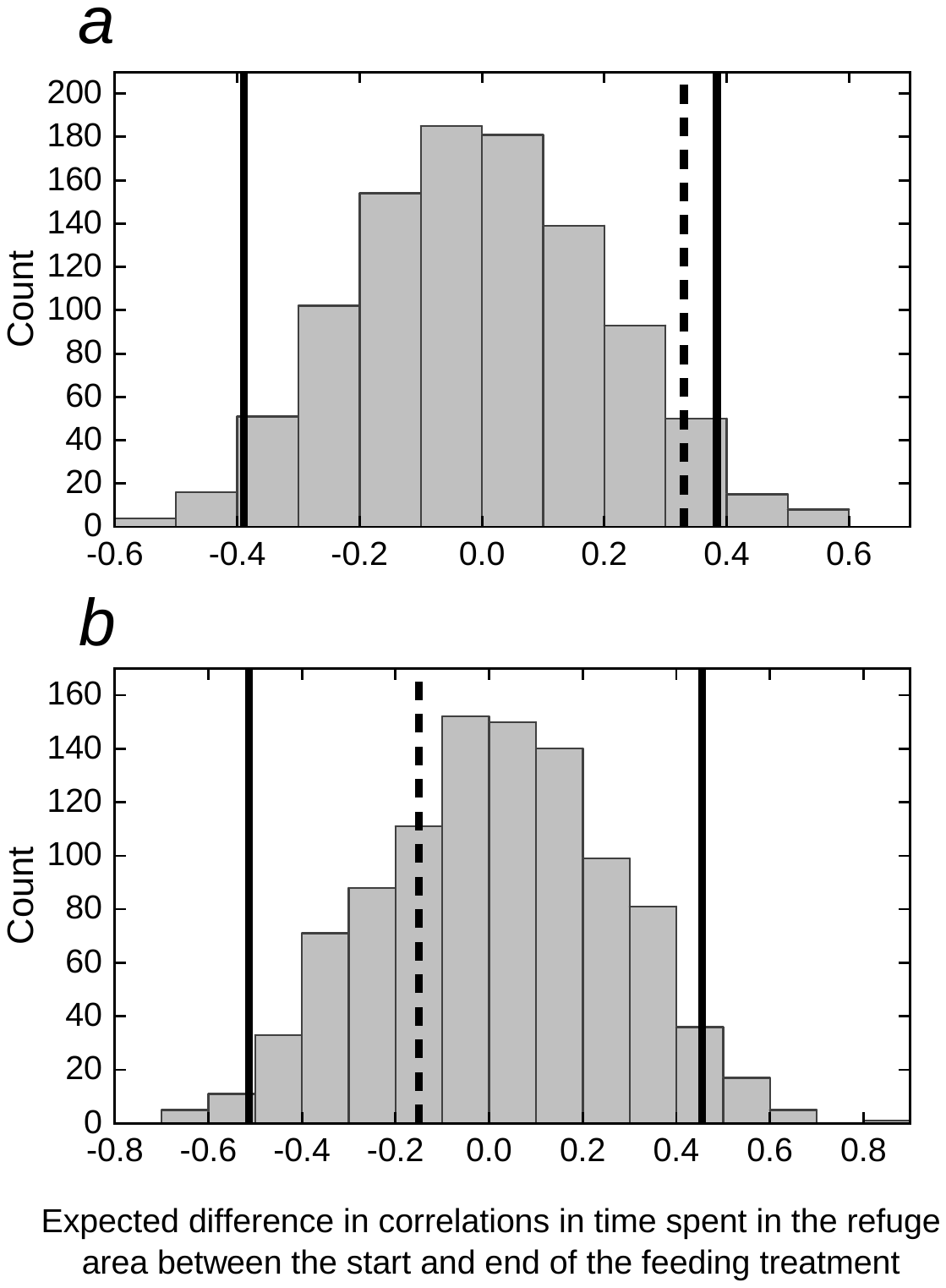

**Figure S3**. The expected difference in Spearman’s rank correlation coefficients in the time spent in the refuge area across the start and across the end of the feeding treatment trials for (a) bold fish and (b) shy fish. Values are based on 1,000 randomizations of the data within individuals. If the observed difference (dashed black line) is outside of the 95% limits of the randomized data’s distribution (solid black lines), it is unlikely that observed difference occurred by chance. Fish were categorized as bold or shy based on their mean latency to emerge from the refuge in control treatment (threshold set as the median value: 246.75 s).

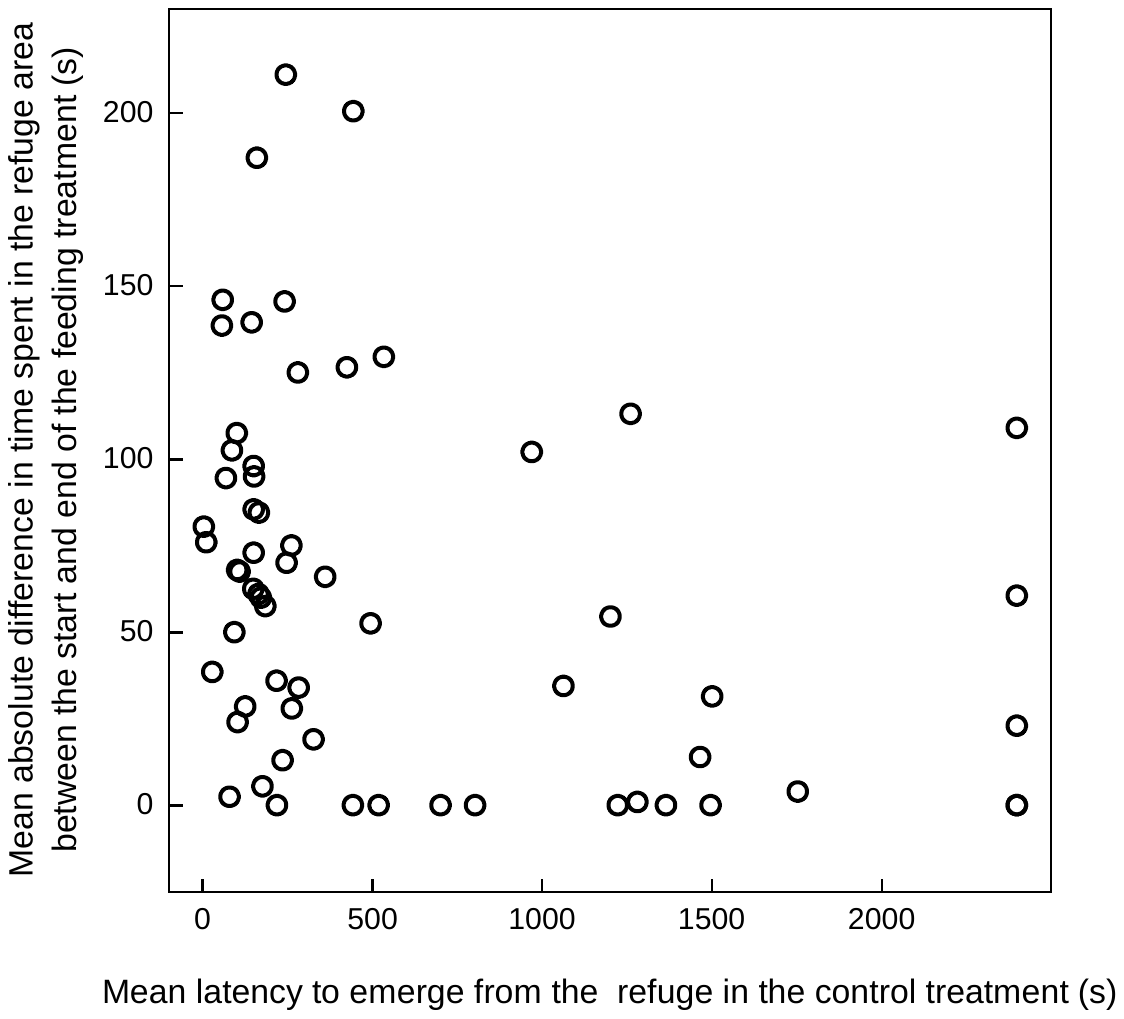

**Figure S4**. The relationship between the mean latency to emerge from the refuge in the control treatment (i.e. boldness) and the mean absolute difference in time that fish spent in the refuge area between the start and end of the feeding treatment.

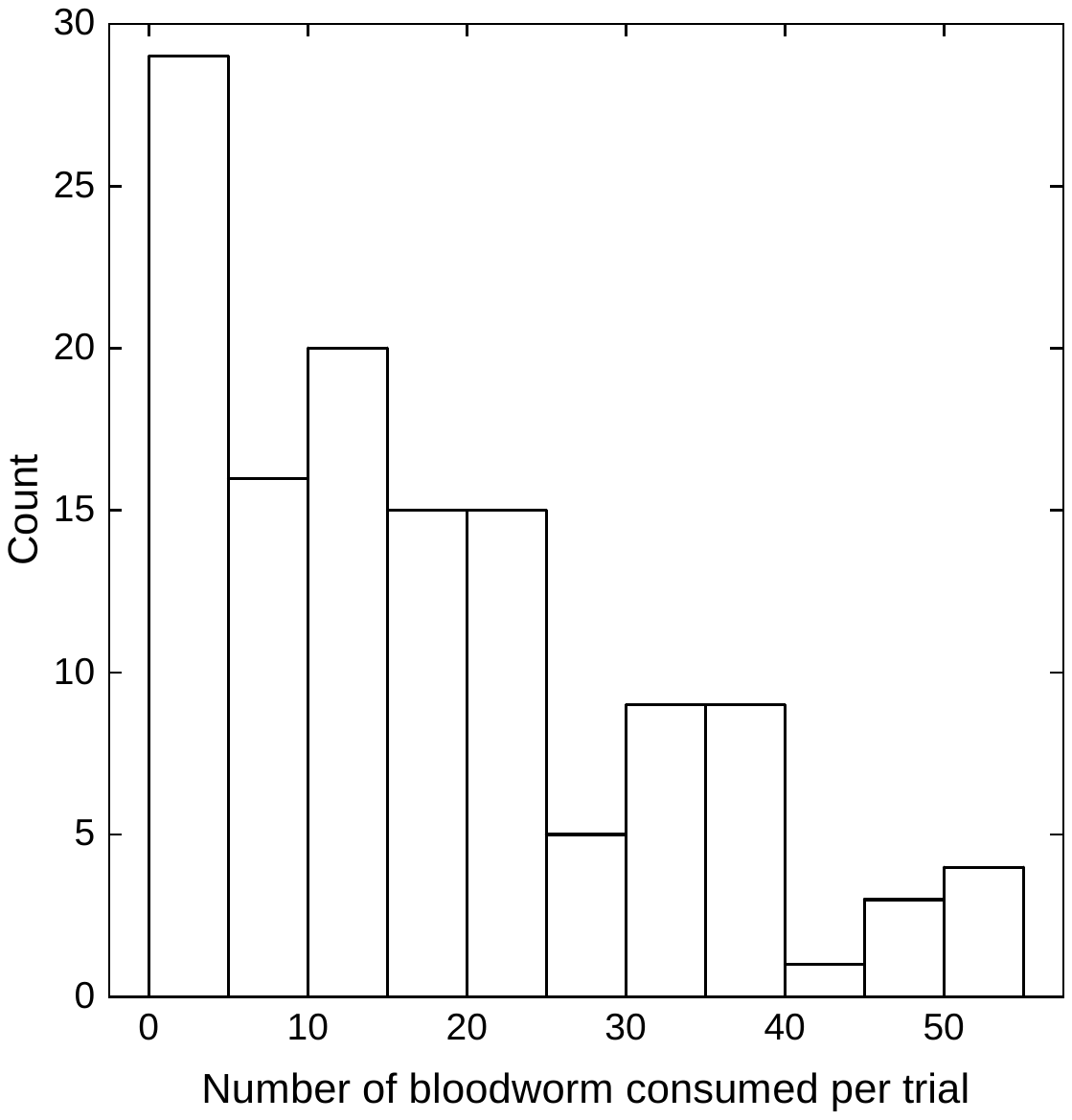

**Figure S5**. The number of bloodworm consumed per feeding treatment trial (n = 126). Fifty bloodworm were available per trial.
